# Integrated surveillance of sandflies as vectors of leishmaniasis reveals the presence of a new *Leishmania* species with possible implication in human health in Europe

**DOI:** 10.1101/2024.05.06.592636

**Authors:** Ignacio Ruiz-Arrondo, Cristina Cervera-Acedo, María Inés Villa-López, Eva María Muelas, Manuel Méndez, José Antonio Oteo, Francisco Collantes

## Abstract

Leishmaniasis, endemic in Spain, displays rising incidence rates, notably in the Region of Murcia. Sandfly surveillance and *Leishmania* screening in 2021 uncovered *Leishmania infantum* and *L. adleri*. While current *Leishmania* diagnostic methods primarily target *L. infantum*, our study underscores the importance of broader *Leishmania* species surveillance for accurate diagnosis and understanding of disease dynamics. The detection of *L. adleri*, previously undescribed in Europe, highlights the need for expanded research to investigate its presence and implications for human and animal health.

---

Leishmaniasis is an endemic zoonosis in Spain, with varying prevalence depending on the geographical area [1]. Since 2015, leishmaniasis has been a notifiable disease in Spain [2] with an incidence rate (IR) of 1 case/100,000 inhabitants. The highest incidence rates in the period 2019-2021 occurred in various regions of the Mediterranean east coast, including the Valencian Community, the Balearic Islands, and the Region of Murcia. In fact, the IR in Murcia is rising faster than the national average [3–5] (Figure 1). Furthermore, the increase in visceral leishmaniasis (VL) in Murcia appears to be associated with apparently healthy individuals, in contrast to the past when it was consistently linked with some form of immunodeficiency [6]. It is essential to develop integrated vector surveillance to better understand the epidemiological changes observed in leishmaniasis. For this reason, a sampling effort of sandflies and potential *Leishmania* carriers was conducted in 2021, near the residences of the leishmaniasis human cases in the Region of Murcia. The protozoan species responsible for both the cutaneous leishmaniasis (CL) and VL in the Iberian Peninsula is *Leishmania infantum*, whose main vector is *Phlebotomus perniciosus* and, to a lesser extent, *P. ariasi* [7]. Both species of sandflies have been reported for decades in the Region of Murcia and neighbouring provinces [8].

**Figure 1:**
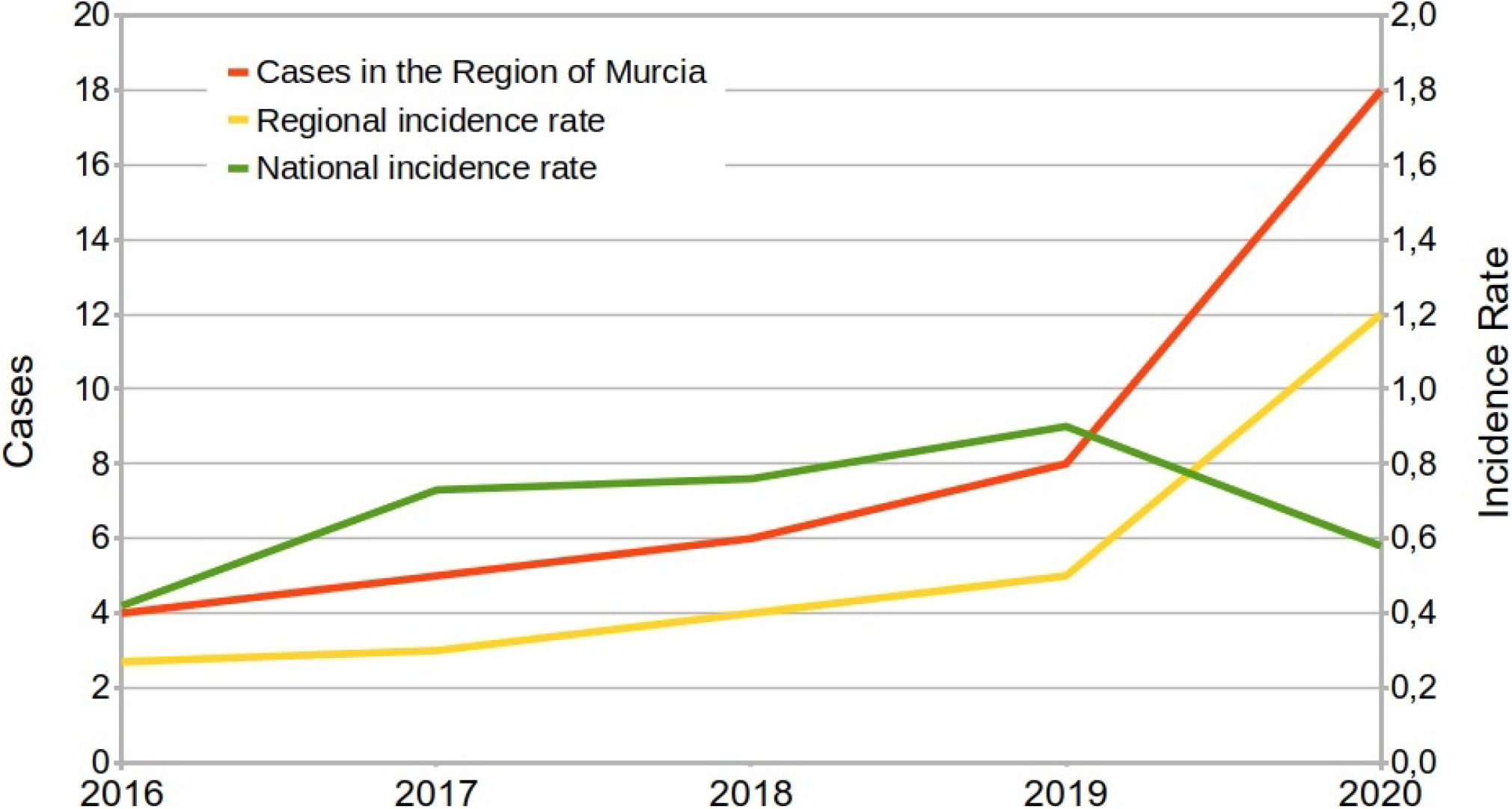
Cases and incidence rate (case/100,000 inhabitants) of leishmaniasis in the Region of Murcia. (southern Spain) [3–5].

Sandfly collection was conducted at 18 sampling points across five municipalities (Figure 2) during June and September 2021, coinciding with the two population peaks of sandflies in the Region of Murcia. Sandflies were collected using paired BG-Pro light traps (LT) baited with CO_2_, one with white light (WL) and other with ultraviolet light (UV), as well as castor oil sticky traps (ST). The LT operated for 24 hours, while the ST remained in place for four days. Samples from the LT were immediately processed on a cold table; Females from LT were morphologically identified based on head and genitalia structures. The rest of the body was preserved in Dulbecco’s Modified Eagle Medium (DMEM) at -80 ºC. ST samples were identified following the same procedure and preserved in absolute ethanol at -80 ºC.

**Figure 2:**
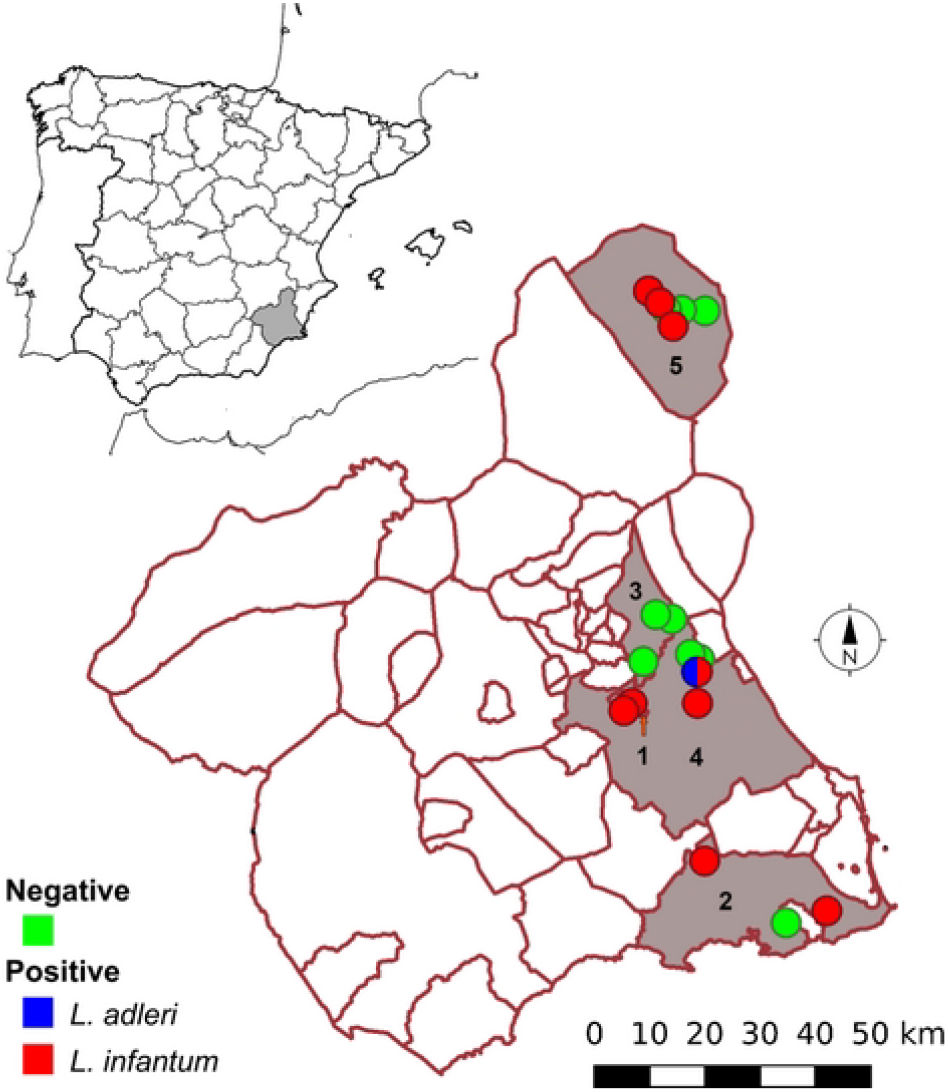
Location of the BG-Pro light traps and results of screening for *Leishmania* spp. in sandflies collected in five municipalities in the Region of Murcia (southern Spain) during June and September 2021. Green circles: sandflies negative for *Leishmania* spp.; red circles: sandflies positive for *Leishmania infantum*; blue circle: sandflies positive for *Leishmania adleri*. Municipalites: 1) Alcantarilla, 2) Cartagena, 3) Molina de Segura, 4) Murcia and 5) Yecla.

Genomic DNA was extracted from individual female sandflies. *Leishmania infantum* specific qPCR was performed, targeting a conserved region in the kinetoplast DNA minicircle [9], with a modification in probe concentration (600 nM). Conventional PCR targeting a partial region of the rRNA internal transcribed spacer 2 (ITS2) was also performed as previously described [10]. In order to better characterize *Leishmania* species different from *L. infantum*, amplification of a larger fragment of the ITS2 was carried out by conventional PCR [11]. Amplicons of the expected size were sequenced using both directions. Nucleotide sequences were compared with those available in GenBank using BLAST (MegaBlast option; https://blast.ncbi.nlm.nih.gov/Blast.cgi).

A total of 468 females sandflies were collected in the study area, with 218 collected in June and 250 in September. Of these 396 were captured by LT and 72 by ST. Morphologic analysis identified three sandfly species (*P. perniciosus, P. papatasi*, and *Sergentomyia minuta*). *Leishmania* DNA was detected in 35 females (7.48%) collected in 4/5 study municipalities, identified as *P. perniciosus* (n=20), *P. papatasi* (n=6) and *S. minuta* (n=9) (Figure 2). *Leishmania infantum* was identified in 34 specimens by qPCR, six of which were also positive through ITS2 (100 % identity with sequences of *L. infantum* MT497972 and MN991197). One *S. minuta* specimen showed a positive result only by ITS2, whose sequence revealed 99.3% identity with *Leishmania* (*Sauroleishmania*) *adleri* from China (HQ289858). A 767 bp fragment of ITS2 was submitted to GenBank under the accession number PP543684.

*Leishamania adleri* was described as a new species from lizards in Kenya in 1958 [12] and our finding constitute the first isolated in sandflies in Europe. This *Leishmania* species is included in the subgenus *Sauroleishmania*, traditionally affecting cold-blooded vertebrates and previously associated with sandflies of the genus *Sergentomyia. Leishmania adleri* is capable of infecting mammals [13], causing transient skin symptoms in humans [14] and inducing asymptomatic infections in hamsters and mice [12]. This species has also been implicated as the likely cause of human leishmaniasis in Ethiopia and China [15,16]. *Sergentomyia minuta*, a species commonly found in our study area, is capable of transmitting *Leishmania tarentolae* [17]. Although *S. minuta* is usually considered to be a cold-blooded feeder, the presence of several specimens with *L. infantum* points to a possible feeding on mammalian hosts, as previously recorded [18]. In addition, recent reports have indicated *L. tarentolae* infections in humans, dogs, and cats [19,20]. These findings suggest the hypothesis that *S. minuta* might potentially be a vector of *L. adleri* in our environment.

The presence of *L. adleri* in the Iberian Peninsula raises several possible hypotheses about its introduction. The first is that it could have been introduced through an infected host, most likely via some species of African reptile for commercial purposes or through a small reptile via sea or air cargo. Another feasible hypothesis could be that it was introduced by an infected person from Sub-Saharan Africa. A less plausible hypothesis of its introduction could be through an infected sandfly in some kind of cargo. These hypotheses provide different starting points for further research on the origin and spread of this species in the study region.

Most studies focused on detecting *Leishmania* spp. in Spain utilize specific targets for detecting *L. infantum* [i.e. 21], as it is the main *Leishmania* species implicated in human and canine cases. According to our results, the ITS2 gene has demonstrated lower sensitivity in detecting *L. infantum* compared to the kinetoplast DNA minicircle. Nonetheless, it has allowed the detection of *L. adleri*, another *Leishmania* species, with potential zoonotic implications. The review of hospital cases of leishmaniasis in Murcia [7] outlines the diagnostic methods employed, including *Leishmania* antibody-ELISA, *Leishmania* antigen in urine and PCR, without specifying the *Leishmania* species detected. Since usual leishmaniasis diagnostic methods involve non-specific tests (i.e. serology) or tests specific for *L. infantum* (i.e. PCR), the results may overlook other *Leishmania* species or assumed to be *L. infantum* by default.

Given the detection of this new species of *Leishmania* in the Iberian Peninsula, further research is warranted to investigate the presence of *L. adleri* in the urban environment on the Region of Murcia and in other areas of the Iberian Peninsula. Its potential presence should be considered in future studies involving sandflies and mammalian and reptile hosts, as well as human samples. The presence of a new *Leishmania* species, previously undescribed in Europe, with potential health implications for humans and other mammals such as pets (dogs and cats) indicates the importance of broadening the spectrum of *Leishmania* species studied to improve the diagnosis of human and pet cases in Spain. Furthermore, new research approaches are needed to better understand the dynamics of leishmaniasis in the Region of Murcia.

## Ethical statement

Ethical approval was not needed for the study.

## Use of artificial intelligence tools

None used.

## Data availability

No applicable

## Authors’ contributions

FC and IR-A drafted the first version of the manuscript. FC provided the maps and the graph. FC, MIV-L, EMM and MM arranged the sampling programme. FC supervised and carried out part of sampling and sandfly identification. IR-A and CC-A performed the molecular analyses. FC, CC-A and IR-A processed the result data. All authors reviewed and contributed to the writing of the manuscript and approved the final version. The views expressed in this manuscript are solely those of the authors and do not necessarily reflect the views, decisions or policies of the institutions with which the authors are affiliated.

